# Decoding functional genes across species with annotation-independent machine learning

**DOI:** 10.1101/2025.07.04.662730

**Authors:** Yanghe Xie, Leyu Li, Geyi Zhu, Peicheng Lu, Jinming Ma, Wenjia Tao, Jinglei Tang, Chuanhong Wang, Guomin Han

**Affiliations:** School of Life Sciences, Anhui Agricultural University, Hefei 230036, China; Institute of Health Sciences and Technology, Anhui University, Hefei 230061, China; National Engineering Laboratory of Crop Stress Resistance Breeding, Anhui Agricultural University, Hefei 230036, China

**Keywords:** Functional Genomics, Machine Learning Framework, Comparative Genomics, Gene Discovery, Complex Traits, Annotation-Independent

## Abstract

Dissecting the genetic basis of complex traits across species is a challenge for traditional low-throughput, species-specific methods. To overcome these limitations, we introduce a computational framework that integrates cross-species comparative genomics and machine learning to identify functional genes for shared phenotypes, independent of prior annotation. Our method links orthologous gene group (OG) profiles to specific traits, enabling the powerful discovery of functional genes across diverse evolutionary clades. For plant–arbuscular mycorrhizal symbiosis, a notoriously difficult system, our model pinpointed a regulatory network by identifying 27 core plant-AM symbiosis genes among its top 50 candidate OGs, including the critical receptor *SYMRK*. The approach also proved highly effective for identifying functional genes related to the tested vertebrate skeletal development and multiple bacterial traits. Most notably, for bacterial motility, our model not only identified 63 known motility genes from the top 100 candidate OGs (of which 78 are present in *Escherichia coli*) but also guided the experimental validation of three novel essential genes. This annotation-independent strategy represents a paradigm shift in functional genomics, offering a scalable and universal engine to decode the genetic architecture of complex traits and illuminate the vast ‘functional dark matter’ across the tree of life.

## Introduction

Elucidating specific gene functions is a paramount objective in modern life sciences, underpinning our comprehension of complex biological systems, the enhancement of economically important traits in agriculture, and the development of therapeutic strategies for human diseases ^1,2^. Current functional gene identification methods primarily rely on forward genetics—associating observed phenotypes with causative genetic loci through mutagenesis or natural variation screening (e.g., Genome-Wide Association Studies, GWAS) ^2–4^—and reverse genetics, which infers gene function by assessing phenotypic alterations resulting from the manipulation of known genes (e.g., via gene knockout or CRISPR-Cas9 editing) ^5–7^. While these traditional methods have proven their utility, they face significant hurdles in dissecting the genetic architecture of complex shared traits—phenotypes conserved across diverse species due to shared ancestral mechanisms. Existing functional gene identification methods are all based on single species; such approaches are often species-specific, labor-intensive, costly, and protracted. For instance, generating comprehensive mutant libraries, even in model organisms, can take years and is complicated by issues like genetic redundancy, lethality, and pleiotropy ^4,8^, while localizing causative mutations from random mutagenesis requires extensive genetic mapping. Similarly, GWAS necessitates large, precisely phenotyped cohorts, often requiring years of material collection or preparation.

Complex shared traits, including developmental programs, stress responses, or morphogenetic processes, such as plant-arbuscular mycorrhizal (AM) fungi symbiosis (regulated by genes like *SYMRK, DMI2, RAM1*), are governed by intricate networks of gene interactions and gene-environment interplay ^9–14^. Elucidating their genetic basis is a formidable task, often requiring sustained, multi-laboratory efforts. Existing methods typically identify only one gene at a time. Consequently, despite the explosion of genomic data, the functions of numerous genes, especially in non-model organisms or for novel roles of known genes, remain uncharacterized, and key functional genes for a vast number of scientifically unexplored traits await discovery ^15,16^. This underscores an urgent need for innovative, high-throughput strategies capable of rapidly and accurately identifying multiple candidate functional genes underlying these complex traits.

Here, we present a cross-species genomics-based computational framework grounded in evolutionary theory: that shared traits across species often arise from functionally conserved orthologous genes, while phenotypic divergence reflects gene specialization or loss. We hypothesized that comparative genomics, coupled with machine learning, could uncover key genes underlying shared traits without relying on prior functional annotations. Our method constructs a feature matrix from orthologous gene group profiles (e.g., presence/absence or copy number) across species and associates these with phenotypic distributions using machine learning. This enables efficient, high-throughput discovery of candidate genes with markedly improved functional localization (Fig. 1). We apply this approach across diverse systems—including plant–arbuscular mycorrhizal symbiosis, vertebrate skeletal traits, and bacterial cell morphology, sporulation, and motility—demonstrating broad utility and robustness. As genome data rapidly accumulate, especially for non-model organisms, this annotation-independent strategy offers a scalable and generalizable tool for elucidating the genetic basis of complex traits across the tree of life.

**Figure 1.**
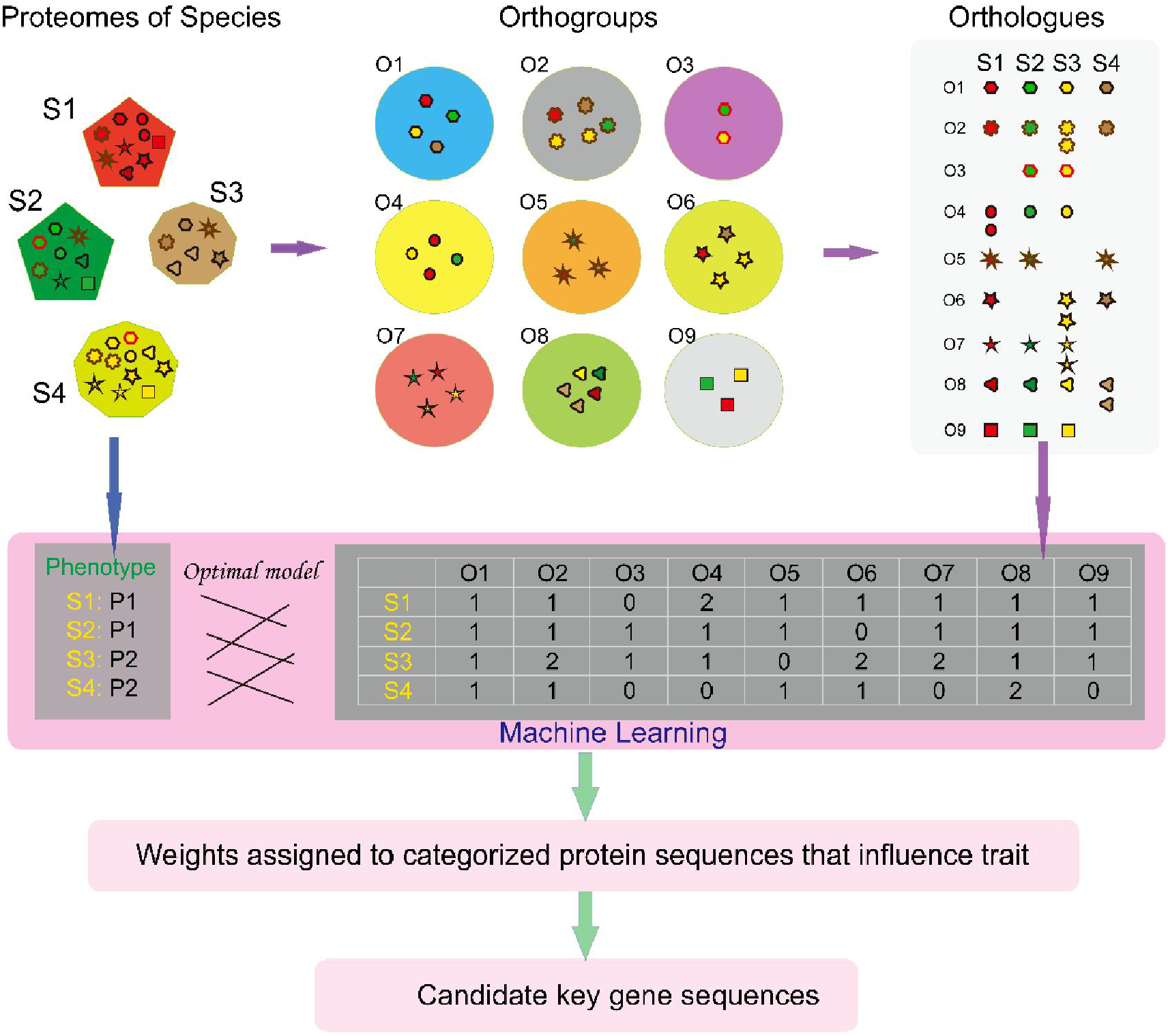
Technical workflow of the annotation-independent gene identification method. Note: All protein sequences encoded by the genomes of various species are compared using the orthology inference software OrthoFinder. A feature matrix is constructed based on the count of sequences assigned by the software to provisional orthologous groups for each species. This matrix, along with phenotypic data, is used to train machine learning models. Accurate models are built, and the contribution of each provisional orthologous group feature to the phenotype is analyzed. Provisional groups with high contribution scores are considered important candidates for identifying key functional genes.

## Results

### Machine learning models link Orthogroup profiles to plant symbiotic phenotypes

OrthoFinder is an excellent tool for inferring orthologous genes based on sequence similarity and evolutionary relationships ^17,18^, widely used in comparative genomics and phylogenetic studies. It performs all-versus-all protein sequence comparisons across multiple species, constructs gene trees based on these alignments, infers orthologous and paralogous relationships, and assigns gene sequences from different plants to distinct provisional orthologous sequence numbers. This entire process operates independently of any third-party alignment or annotation databases. Leveraging this characteristic of OrthoFinder, we attempted to associate matrices of these provisional orthologous sequences with phenotypes using various machine learning models. We first investigated plant-AM fungi symbiosis, a complex shared trait. Using OrthoFinder, we clustered proteomes of 141 plant species (68 symbiotic, 73 non-symbiotic, Supplementary Table 1) and constructed a feature matrix based on OG gene counts per species. Five machine learning algorithms were tested. Decision Tree (DT) and Conditional Inference Tree (CTree) models achieved the highest performance on the test set (accuracy >95%, precision >96%, recall >94%, AUC 0.96; Fig. 2A-B). Random Forest (RF) also performed robustly (accuracy 94.29%, AUC 0.94; Fig. 2a-b). This demonstrates that OG profiles effectively capture genomic signatures distinguishing symbiotic capability.

**Figure 2.**
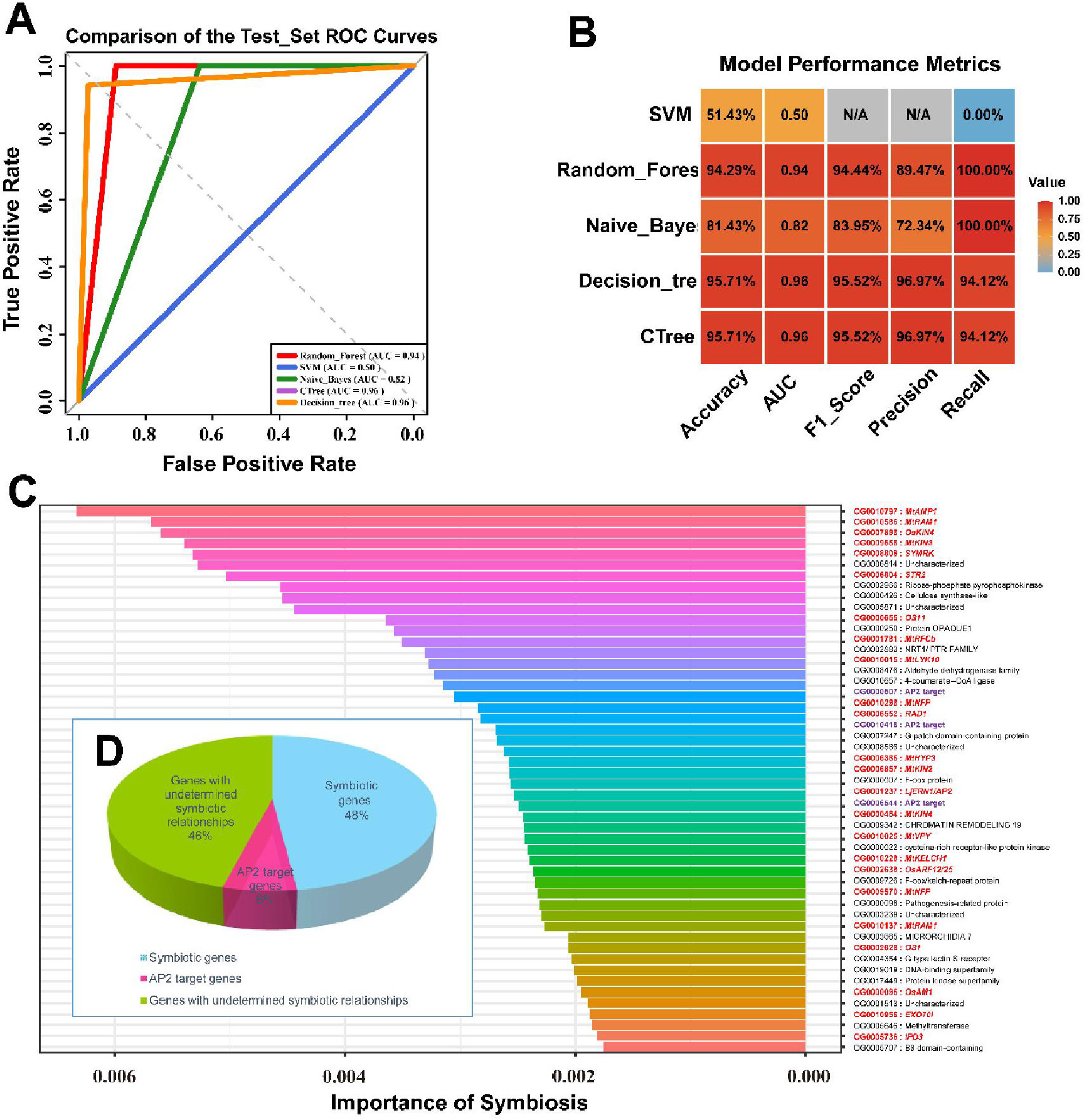
Evaluation of machine learning models for plant-AM fungi symbiosis and functional analysis of important candidate sequences. (A) ROC curves for various machine learning algorithms on the test set for plant-AM fungi symbiosis classification: CRTree, Conditional Inference Tree; Decision_tree, Decision Tree; SVM,Support Vector Machine; Naive_Bayes, Naive Bayes; Random_Forest, Random Forest. (B) Heatmap of performance metrics (Accuracy, AUC, F1 Score, Precision, Recall) for multiple algorithms. (C) Bar chart of the top 50 features ranked by the Random Forest “Symbiosis” importance metric, with annotations: Red font indicates known AM symbiosis related genes; purple font indicates AP2 downstream target genes; black font indicates genes whose relationship to the symbiosis pathway is currently undetermined. (D) Pie chart summarizing the analysis of the top 50 most important sequences identified by the novel method, showing that 54% are key known genes directly related to symbiosis.

### The OrthoFinder-Random Forest model identifies known and novel symbiosis genes

The RF model’s feature importance ranking highlighted OGs crucial for predicting symbiotic phenotypes. Functional annotation of the top 50 most important OGs revealed containing orthologs of known core plant-AM symbiosis genes (e.g., *SYMRK, MtNFP, MtKIN2/3, OsKIN4, IPD3, MtRAM1, RAD1, EXO70I, STR2, MtAMP1,MtRFCb, MtHYP3, LjERN1*), as well as three symbiosis-related AP2 transcription factor targets. These genes are involved in signal transduction, symbiotic structure development, nutrient exchange, and regulation (Fig. 2C-D, Supplementary Table 2). This efficient identification of numerous validated key genes, directly from sequence-derived OG profiles and phenotype labels, represents a significant advance.

To assess the impact of transcript diversity on model performance and feature selection, we constructed a “unique protein model” by retaining only one representative transcript per gene locus. This model also accurately predicted symbiotic capability based on genomic features (Supplementary Figure 1). Notably, while yielding comparable predictive performance to the all-transcripts model, the unique protein model enhanced the discovery of known symbiosis genes within broader top-ranked lists (e.g., identifying 51 vs. 45 known genes in the Top 200 OGs, respectively; Fig. 3A), suggesting improved stability and potentially greater discovery power for this refined feature set. We further evaluated different RF importance metrics. The “Symbiosis” metric (based on class prediction), “MeanDecreaseAccuracy,” and “MeanDecreaseGini” all yielded largely concordant feature rankings, though “MeanDecreaseGini” identified slightly more known symbiosis-related genes (Fig. 3B). Many other highly-ranked OGs without prior functional links to symbiosis exhibited similar conservation patterns—highly conserved in symbiotic plants and exhibiting low conservation or absence in non-symbiotic plants—suggesting they are strong candidates for novel symbiosis genes (Fig. 3C, Supplementary Table 3). Model stability was confirmed by averaging results from 10 runs with different random seeds; key symbiosis genes consistently maintained high rankings (Supplementary Table 4 & Supplementary Table 5).

**Figure 3.**
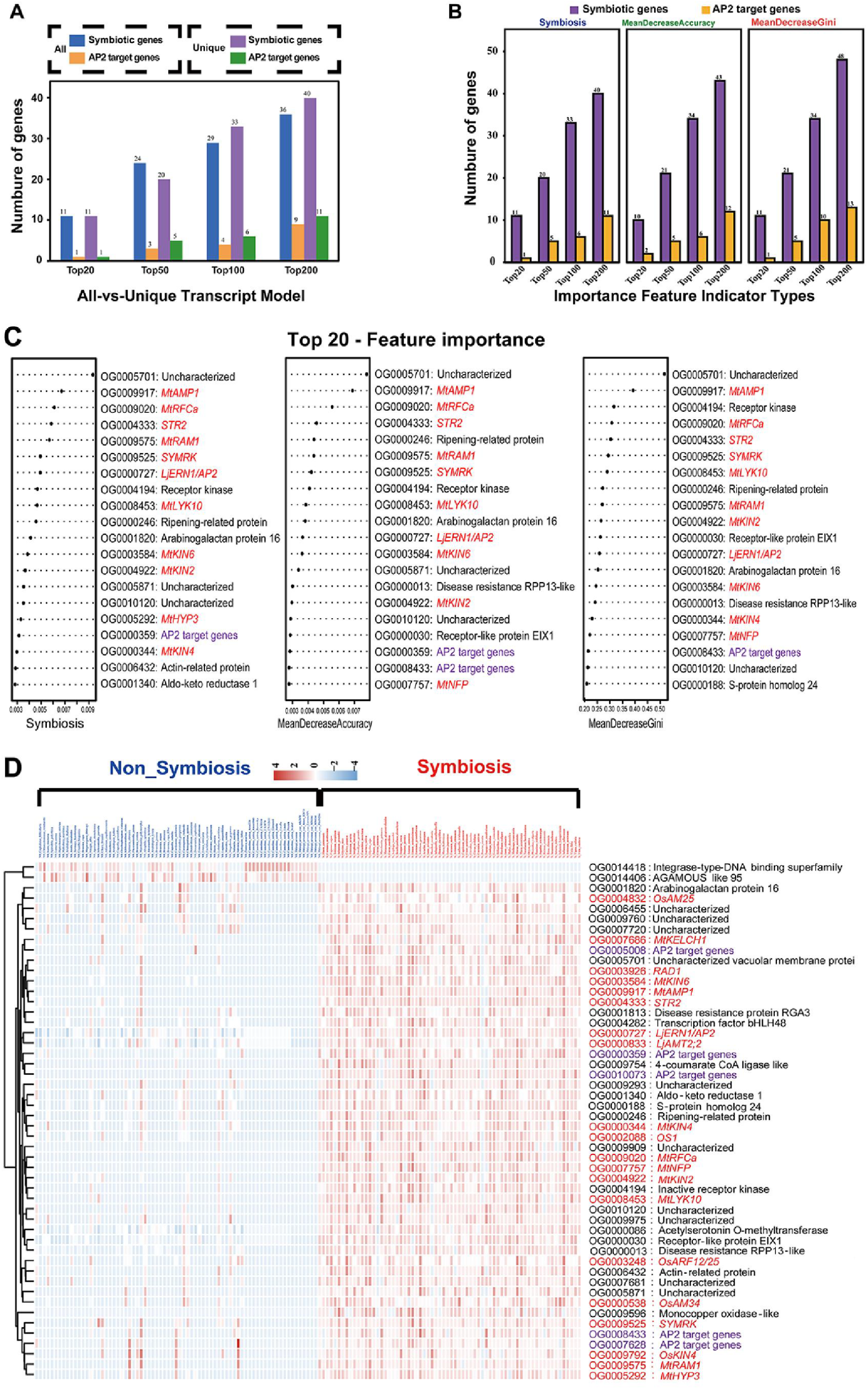
Comparison of machine learning model performance optimization and distribution patterns of candidate sequence features in symbiotic versus non-symbiotic plants. (A) Comparison of the number of known symbiosis genes among the top 20, top 50, top 100, and top 200 candidate sequences identified by the all-transcripts model versus the unique-transcript model. Blue and orange bars represent the number of known symbiosis genes and AP2 downstream target genes, respectively, identified by the all-transcripts model. Purple and green bars represent the number of known symbiosis genes and AP2 downstream target genes, respectively, identified by the unique-transcript model. (B) Bar chart comparing the number of known symbiosis genes (purple) and AP2 downstream target genes (orange) within the top 20, top 50, top 100, and top 200 ranked features using three Random Forest importance metrics: “Symbiosis”, “MeanDecreaseAccuracy”, and “MeanDecreaseGini”. (C) Specific ranking order within the top 20 features using the “Symbiosis”, “MeanDecreaseAccuracy”, and “MeanDecreaseGini” importance metrics. Red and purple indicate known symbiosis genes and AP2 downstream target genes, respectively. (D) Distribution patterns of candidate sequence feature values (derived from the unique-transcript prediction model) in symbiotic versus non-symbiotic plants. Red and purple indicate known symbiosis genes and AP2 downstream target genes, respectively.

### Universality demonstrated in vertebrates and bacteria

To test broader applicability, we applied the method to 409 animal genomes (189 vertebrates, 220 invertebrates). RF and DT models achieved highly accurate classification (accuracy >99%, AUC ∼1.0; Supplementary Table 6, Supplementary Figure 2A-B). Among the top 50 OGs, 34 were associated with vertebrate-specific traits, including 18 potentially involved in skeletal development and diseases (e.g., *Mfap2, Map3k14, Acod1, Bend3, FAM83H, Bri3bp, Lbh, P2rx7*), 9 in neurodevelopment, 4 in haematopoiesis, and 3 in muscle development (Supplementary Table 7), many of which are highly conserved or specific to vertebrates.

The strategy was then applied to prokaryotes. For bacterial rod-shape determination (1330 rod-shaped, 1140 coccoid bacteria; Supplementary Table 8), the RF model achieved 95.12% accuracy (AUC 0.95; Supplementary Figure 2C-D). Top OGs included known rod-shape determinants like *pal* (OG0000056) and *mreB* (OG0000186), cell wall remodeling gene *nlpD* (OG0000064), and cell division regulators *ftsL* and *dapF* (Supplementary Table 9).

For bacterial spore formation (221 spore-forming, 220 non-spore-forming; Supplementary Table 10), the RF model achieved 94.47% accuracy (AUC 0.95; Supplementary Figure 2E-F). The top 50 OGs included 12 known key sporulation/germination genes like *sigB* (OG0000159), *aprE* (OG0000063), *divIVA* (OG0000539), *spoVK* (OG0001150), and *gerKA* (OG0000442) (Supplementary Table 11).

### Experimental validation of novel motility genes in *Escherichia coli*

We further challenged our method to identify bacterial motility genes (487 motile, 487 non-motile species, Supplementary Table 12). The RF model performed best (accuracy 85.17%, AUC 0.85; Fig. 4A-B). Among the top 100 OGs predictive of motility, of which 78 were found in the genome of *E. coli* AE81, 63 contained experimentally verified motility or chemotaxis genes (Table 1). To validate predictions for novel candidates, we focused on OGs within the top 100 that were uncharacterized for motility in *E. coli* AE81 (Supplementary Table 13). We successfully knocked out five such genes: *pal, yicC, dusA, hslU*, and *glyQ* (Supplementary Figure 3). Growth curves were unaffected (Fig. 4D). Motility assays revealed significantly reduced swimming halo diameters for *Δpal, ΔdusA*, and *ΔyicC* strains compared to wild-type (Fig. 4C&E), confirming their essential role in *E. coli* AE81 motility. This successful identification and experimental validation of novel functional genes underscores the robust predictive power of our annotation-independent approach.

**Figure 4.**
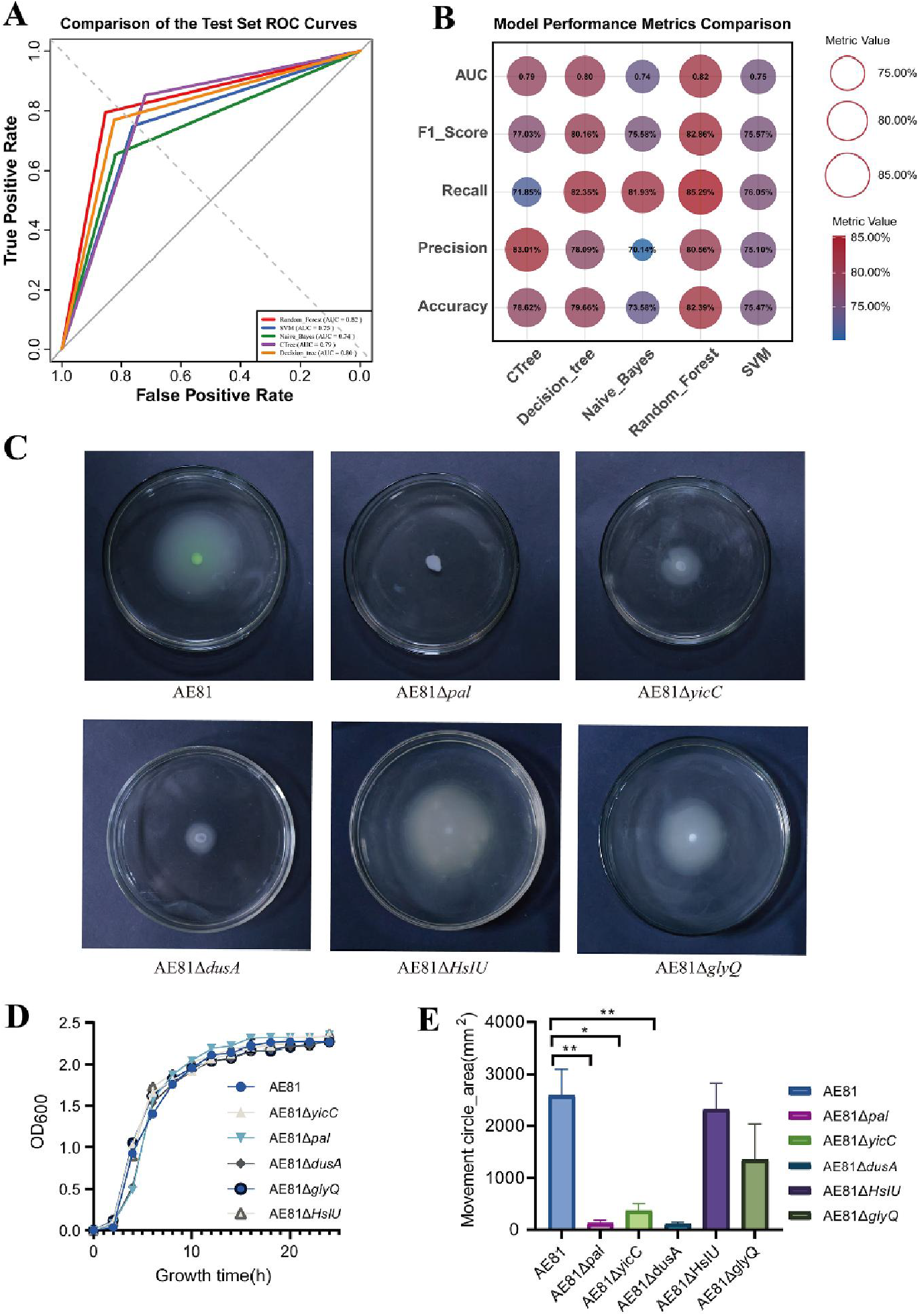
Evaluation of the bacterial motility prediction model and experimental validation of candidate genes. (A) ROC curves for various machine learning algorithms on the test set for bacterial motility classification: CRTree, Conditional Inference Tree; Decision_tree, Decision Tree; SVM, Support Vector Machine; Naive_Bayes, Naive Bayes; Random_Forest, Random Forest. (B) Heatmap of performance metrics (Accuracy, AUC, F1 Score, Precision, Recall) for multiple algorithms. (C) Motility assays following knockout of *pal, yicC, dusA, hslU*, and *glyQ*. Knockout of *pal, dusA*, and *yicC* significantly reduced the motility of *E. coli* AE81. (D) Growth curves of wild-type *E. coli* AE81 and knockout strains, showing that the growth rates of all knockout strains were comparable to the wild-type. (E) Comparison of swimming halo diameters of wild-type AE81 and knockout strains. Statistical significance was determined by F-test: ** indicates *p* < 0.01; * indicates *p* < 0.05.

**Table 1.** Provisional names and functional annotations of the top 50 candidate OG sequences from the bacterial motility prediction model.

| Rank | Orthogroup | Annotation | Rank | Orthogroup | Annotation |
| --- | --- | --- | --- | --- | --- |
| 1 | OG0001106 | *motA* | 26 | OG0001460 | *fliL* |
| 2 | OG0000334 | *flgF* | 27 | OG0000868 | *cheA* |
| 3 | OG0001617 | *fliM* | 28 | OG0002383 | - |
| 4 | OG0001457 | *flgB* | 29 | OG0001665 | *queF* |
| 5 | OG0001443 | *fliG* | 30 | OG0001686 | *yaiI* |
| 6 | OG0001367 | *fliQ* | 31 | OG0000214 | *flhF* |
| 7 | OG0001467 | *fliE* | 32 | OG0000050 | *pal* |
| 8 | OG0000406 | *fljL* | 33 | OG0000552 | *yjcC* |
| 9 | OG0001369 | *fliF* | 34 | OG0001905 | *flgA* |
| 10 | OG0001366 | *fliP* | 35 | OG0001538 | *hfq* |
| 11 | OG0001357 | *flhA* | 36 | OG0000970 | *yuxH* |
| 12 | OG0001453 | *flgC* | 37 | OG0001806 | *flgH* |
| 13 | OG0001574 | *fliK* | 38 | OG0001858 | *flgI* |
| 14 | OG0000950 | *flhB* | 39 | OG0000052 | *acrB* |
| 15 | OG0000997 | *fliN* | 40 | OG0001937 | - |
| 16 | OG0001451 | *flgK* | 41 | OG0000032 | *glyQ* |
| 17 | OG0000021 | *tsr* | 42 | OG0001438 | *dsbB* |
| 18 | OG0001368 | *fliR* | 43 | OG0001478 | *yggT* |
| 19 | OG0001435 | *flgD* | 44 | OG0002355 | *cheZ* |
| 20 | OG0001674 | *fliS* | 45 | OG0001544 | *zapA* |
| 21 | OG0001662 | *fliD* | 46 | OG0001654 | *queC* |
| 22 | OG0000432 | *cheW* | 47 | OG0001873 | *lptD* |
| 23 | OG0001900 | *flgM* | 48 | OG0002251 | - |
| 24 | OG0001690 | *fliH* | 49 | OG0001441 | *hslU* |
| 25 | OG0000302 | *fliI* | 50 | OG0002246 | *prkA* |

Note: Red font indicates known motility-related genes; green font indicates genes with unknown relevance to motility; green font with a yellow background highlights genes successfully knocked out in this study. The symbol "-" represents the absence of the gene in the *E. coli* AE81 genome.

## Discussion

The rapid and accurate identification of critical functional genes responsible for complex biological traits from tens of thousands of candidates poses a significant technical challenge and remains a central hurdle in biology. Traditional methods are often limited in scope, throughput, and applicability to non-model organisms. Our study introduces a powerful computational strategy that directly links gene sequences (via Orthologous Group profiles) to phenotypes across diverse species, representing a significant advance by operating independently of pre-existing functional annotation databases for the initial discovery phase. Existing methods that directly compare sequences against known functional gene sequences in databases like NCBI ^19^ or UniProt ^20^, while capable of analyzing the potential functions of some known genes, often fail to directly associate them with specific phenotypes. Functional classification of species’ sequences based on gene families also presents challenges in directly linking specific family members to particular traits ^21,22^, primarily due to the large number of members within these families. Furthermore, existing annotation methods based on third-party database comparisons are unable to analyze the potential functions of completely novel sequences ^23,24^. Our method contrasts sharply with those that rely on known gene functions to guide analysis or interpret results from the outset.

The core breakthrough of our method lies in providing a novel, high-throughput approach to functional gene identification. It transcends the reliance of traditional methods on random mutations within a single species or natural variations within a population; instead, it leverages the wealth of genomic and phenotypic information accumulated during long-term cross-species evolution. The underlying principle is that shared phenotypes often rely on a conserved set of OGs ^25^, while phenotypic differences correlate with the evolutionary dynamics of these OGs. By leveraging machine learning ^26^, we can systematically mine these patterns from large-scale comparative genomic data. The successful identification of numerous known key genes for AM symbiosis (27 of top 50 OGs, including critical receptors), vertebrate skeletal development skeletal development and bone-related diseases (18 of top 50 OGs), bacterial cell shape (e.g., *mreB, pal*), sporulation and germination (e.g., *sigB, spoVK*), and motility (63 of top 78 OGs in *E. coli* AE81) (Table 1) across vastly different life forms robustly validates the approach. By analyzing the distribution patterns and evolutionary trajectories of OGs across different phenotypic groups, and harnessing the powerful pattern recognition capabilities of machine learning, our study achieves a technological leap in rapidly and systematically predicting potential key functional gene groups directly from macroscopic, shared phenotypic differences, with an identification efficiency far exceeding existing gene discovery methods.

For the diverse traits tested, our method identified in a single analysis nearly as many core genes as traditional methods have identified over decades of research, and additionally flagged many strong novel candidates based on their evolutionary patterns and model importance. Regarding genes related to animal skeletal development, our method not only identified genes involved in early skeletal development (e.g., mutations in *Bri3bp* are associated with bone deformity ^27^, and *Lbh*, a highly conserved transcription factor, may participate in endochondral ossification ^28^) but also many genes related to skeletal diseases and potential new therapeutic targets. For instance, *P2rx7* plays a key role in osteocyte formation and function and has become a potential target for osteoporosis treatment ^29–31^; the osteosarcoma-related gene *Cdc42ep3* was also identified ^32^, suggesting utility in understanding complex developmental and pathological processes. Notably, plant receptor gene (*SYMRK*) crucial for recognizing AM fungi symbiosis ^33,34^ was also identified by our method. The precise identification of plant receptors involved in microbial interactions is notoriously challenging; for instance, plant receptors for mycorrhizal symbiosis were only identified following the isolation and characterization of fungal Myc factors ^35^. The direct identification of such receptor gene sequences among the top candidates by our novel approach underscores its remarkable power.

The broad applicability of this strategy is a key strength. Its success in plants, animals, and bacteria for traits ranging from symbiosis to morphogenesis and organ development demonstrates its potential as a universal tool. This is particularly valuable for exploring the vast “functional dark matter” in non-model organisms or for uncovering novel functions of known genes. The experimental validation of *pal, dusA*, and *yicC* as novel motility genes in *E. coli*, identified from relatively lower rankings (within top 100), further attests to the method’s reliability in generating testable hypotheses and discovering new biology.

We meticulously evaluated model stability and parameter choices. Several distinct feature importance metrics within RF algorithm effectively identified key functional genes. Among these, the “MeanDecreaseGini” ^36^ metric generally provided the most comprehensive identification of known symbiosis genes, while metrics reflecting specificity (e.g., class-specific importance for “Vertebrate”) performed better in the vertebrate analysis. The optimal metric may thus vary depending on the specific research context and requires empirical comparison. Further analysis revealed that these metrics are better suited for distinguishing between different types of predictive features; some features are highly prevalent in one phenotypic group and have low frequency in another. Therefore, combining ML output with evolutionary conservation patterns (e.g., high prevalence in phenotype-positive species, low prevalence or absence in phenotype-negative species) makes functional gene identification more reliable, helps reduce the workload of subsequent experimental validation, and provides a robust dual-filter criterion. Although machine learning is a “black-box” model and results might vary ^37^, using a fixed seed ensures identical results on different computers. However, changing the seed value can lead to some differences ^38^. We tested ten runs with different random seeds, and the results showed that model outputs remained stable across multiple runs. While there were slight variations, the top-ranking major results generally remained at the top, ensuring reproducibility.

Despite its successes, the method has limitations. Some known key genes (e.g., *CCaMK* in plants ^39^, *Spo0A* in bacteria ^40^) were not always top-ranked. This may reflect factors such as suboptimal genome assembly/annotation quality in some input species, biases in species sampling, limitations of OG clustering algorithms with complex evolutionary events (gene fusion/fission, HGT), insufficiently refined phenotype classification (e.g., bacterial motility could be divided into more specific types ^41^), or the specific way certain genes contribute to a phenotype that is not well captured by simple presence/absence/copy number in the sampled species. The “black-box” nature of some ML models can also make interpretation challenging, and the fixed number of top features analyzed is a simplification. Future advancements in genome sequencing (e.g., T2T assemblies), broader and more phylogenetically balanced species sampling, improved OG inference algorithms, and more interpretable machine learning models will undoubtedly enhance the power and precision of this approach.

In conclusion, we have developed and validated a novel, annotation-independent computational strategy that directly links gene sequence repertoires to complex shared phenotypes across the tree of life. This represents a paradigm shift from reliance on single-species studies or existing functional databases for discovery, offering a scalable and efficient engine for functional genomics. This new method promises to accelerate the dissection of the genetic architecture of complex biological traits, illuminate evolutionary mechanisms, and provide a powerful research tool for understanding biological mysteries and developing therapeutic interventions.

## Methods

### Genome Data Acquisition

Plant genomes (141 species: 68 symbiotic, 73 non-symbiotic with AM fungi) were sourced from NCBI, GWH, and Phytozome, based on Radhakrishnan et al. ^42^ and public databases (Supplementary Table 1). Animal genomes (189 vertebrates, 220 invertebrates) were from NCBI (Supplementary Table 6). Bacterial genomes for shape analysis (1330 rod, 1140 coccus) (Supplementary Table 8), spore formation (221 forming, 220 non-forming) (Supplementary Table 10), and motility (487 motile, 487 non-motile) (Supplementary Table 12) were downloaded from NCBI ^19^, with phenotypes cross-referenced from BacDive ^43^.

### Orthologous Group Identification and Feature Matrix Construction

OrthoFinder (v3.0.1b1) ^18^ was used with default parameters (DIAMOND for alignment) to identify OGs from whole-genome protein sequences for each dataset. Custom Python scripts were used for sequence renaming. The primary feature matrix for machine learning comprised the count of genes from each species belonging to each OG. An alternative “unique protein model” used only the longest transcript per gene locus. The structure is exemplified in Supplementary Table 14.

### Machine Learning Workflow

Datasets were split into training and test sets (typically 1:1). Five algorithms were evaluated: Random Forest (RF; randomForest R package), Support Vector Machine (SVM; e1071 R package), Naive Bayes (NB; e1071 R package), Conditional Inference Tree (CTree; party R package), and Decision Tree (DT; rpart R package) ^44,45^. Model performance was assessed using Accuracy, Precision, Recall, F1 Score, and Area Under the ROC Curve (AUC) on the test set.

### Identification and Functional Annotation of Important Genes

Feature importance scores from the best-performing models (often RF’s MeanDecreaseGini) were used to rank OGs. For annotation, protein sequences from representative species within top-ranked OGs were extracted. These were compared against curated lists of known functional genes using BLASTp (E-value ≤ 1e-5, identity >30%, query coverage >50%). Unannotated OGs were analyzed using NCBI BLASTp against the non-redundant protein database. Crucially, this annotation step is for interpretation after ML-based identification and does not inform the initial model training or feature selection, ensuring discovery independence.

### Model Optimization and Stability

For plant symbiosis, a “unique protein model” using only the longest transcript per gene was tested. Different RF importance metrics (MeanDecreaseAccuracy, MeanDecreaseGini, class-specific importance “Symbiosis”) were compared. Model stability was assessed by running RF 10 times with different random seeds and averaging feature importance scores.

### Bacterial Motility Gene Knockout and Phenotyping

Candidate motility genes identified in *E. coli* AE81 (APEC strain, gift from Prof. Kezong Qi) were knocked out using a CRISPR/Cpf1 dual-plasmid system ^46^. Briefly, crRNAs targeting genes (PAM: TTV) were cloned into pcrE, followed by 500 bp upstream/downstream homologous arms for each target gene, generating pcrEG-XXX knockout plasmids (primers in Supplementary Tables 15, Supplementary Tables 16). Plasmids were transformed into *E. coli* AE81 harboring pEcCpf1. Knockouts were verified by colony PCR and Sanger sequencing.

For motility assays, overnight cultures were subcultured to OD_600_=1.0, washed, resuspended in PBS to OD_600_=2.0. 2 µL was spotted onto semi-solid LB agar (0.25% agar) and incubated at 37°C for 12h. Swimming halo diameters were measured.

Growth curves were determined by inoculating LB medium (with appropriate antibiotics) with 1:100 revived cultures, grown to mid-log phase (OD_600_ 0.5-0.6), then standardizing and inoculating 100 ml fresh LB at 1% (v/v). OD600 was measured every 2h at 37°C with shaking.

## Acknowledgements

This work was supported by grants from the Anhui Provincial Peak Discipline Project of Biology.

## Supplementary Figure and Table Captions

**Supplementary Figure 1 Evaluation of the unique-transcript machine learning model for plant-AM fungi symbiosis.**

(A) ROC curves for various machine learning algorithms on the test set for plant-AM fungi symbiosis classification using the unique-transcript model: CRTree, Conditional Inference Tree; Decision_tree, Decision Tree; SVM, Support Vector Machine; Naive_Bayes, Naive Bayes; Random_Forest, Random Forest. (B) Heatmap of performance metrics (Accuracy, AUC, F1 Score, Precision, Recall) for multiple algorithms using the unique-transcript model.

**Supplementary Figure 2 Evaluation of prediction models for vertebrate status, bacterial rod shape, and bacterial spore formation.**

(A), (C), (E) ROC curves for various machine learning algorithms on the respective test sets: CRTree, Conditional Inference Tree; Decision_tree, Decision Tree; SVM, Support Vector Machine; Naive_Bayes, Naive Bayes; Random_Forest, Random Forest. (B), (D), (F) Heatmaps of performance metrics for multiple algorithms on the respective test sets. (A), (B) Vertebrate status prediction model. (C), (D) Bacterial rod shape prediction model. (E), (F) Bacterial spore formation prediction model.

**Supplementary Figure 3 Sequencing verification results for each knockout strain.**

**Supplementary Table 1 Species composition and genome source information for AM fungal symbiotic plants.**

**Supplementary Table 2 Provisional names and functional annotations of the top 50 candidate OG sequences from the mycorrhizal symbiosis model.**

**Supplementary Table 3 Distribution of the top 50 candidate OG sequences in the genomes of symbiotic and non-symbiotic plants.**

**Supplementary Table 4 Ranking changes of the top 50 candidate OGs from 10 runs of the OrthoFinder unique-protein model with random seeds.**

Note: Red indicates OGs corresponding to known symbiosis genes; Purple indicates OGs corresponding to AP2 target genes.

**Supplementary Table 5 The top 50 important features and their average ranks from 10 runs of the model with a random seed.**

**Supplementary Table 6 Animal species information used for the vertebrate status prediction model.**

**Supplementary Table 7 Provisional names and functional annotations of the top 50 candidate OG sequences from the vertebrate status prediction model.**

Note: Blue font indicates genes related to skeletal development and bone-related diseases; red font indicates genes related to neurodevelopment and neurological diseases; purple font indicates genes related to hematopoiesis or blood; green font indicates genes related to muscle development.

**Supplementary Table 8 Bacterial species composition information used for the bacterial rod shape prediction model.**

**Supplementary Table 9 Provisional names and functional annotations of the top 50 candidate OG sequences from the bacterial rod shape prediction model.**

**Supplementary Table 10 Bacterial species composition information used for the bacterial spore formation prediction model.**

**Supplementary Table 11 Provisional names and functional annotations of the top 50 candidate OG sequences from the bacterial spore formation prediction model.**

**Supplementary Table 12 Bacterial species composition information used for the bacterial motility prediction model.**

**Supplementary Table 13 Provisional names and functional annotations of the top 100 candidate OG sequences from the bacterial motility prediction model.**

Note: Red font indicates known motility-related genes; green font indicates genes with unknown relevance to motility; green font with a yellow background highlights genes successfully knocked out in this study.

**Supplementary Table 14 Diagrammatic representation of the feature dataset structure. Supplementary Table 15 crRNA synthesis primer sequences.**

**Supplementary Table 16 Homologous arm synthesis primer sequences.**

## Extanded Data Table Captions

**Extended Data Table 1 All feature importance values from the all-transcripts model for plant-AM symbiosis gene identification.**

**Extended Data Table 2 All feature importance values from the unique-transcript model for plant-AM symbiosis gene identification.**

**Extended Data Table 3 All feature importance values from the unique-transcript model for animal skeletal gene identification.**

**Extended Data Table 4 All feature importance values from the model for bacterial rod-shape gene identification.**

**Extended Data Table 5 All feature importance values from the model for bacterial sporulation gene identification.**

**Extended Data Table 6 All feature importance values from the model for bacterial motility gene identification.**

## References

1. Yu, P. et al. Seedling root system adaptation to water availability during maize domestication and global expansion. Nat Genet 56, 1245–1256 (2024).

2. Pairo-Castineira, E. et al. GWAS and meta-analysis identifies 49 genetic variants underlying critical COVID-19. Nature 617, 764–768 (2023).

3. Tian, T. et al. Genome assembly and genetic dissection of a prominent drought-resistant maize germplasm. Nat Genet 55, 496–506 (2023).

4. Xiong, H. et al. A large-scale whole-exome sequencing mutant resource for functional genomics in wheat. Plant Biotechnol J 21, 2047–2056 (2023).

5. Shalem, O. et al. Genome-Scale CRISPR-Cas9 Knockout Screening in Human Cells. Science 343, 84–87 (2014).

6. Wang, L. et al. CRISPR/Cas9-based editing of NF-YC4 promoters yields high-protein rice and soybean. New Phytol 245, 2103–2116 (2025).

7. Aklilu, E. Review on forward and reverse genetics in plant breeding. All Life (2021).

8. Lu, X. et al. Gene-Indexed Mutations in Maize. Mol Plant 11, 496–504 (2018).

9. Shi, J. et al. A phosphate starvation response-centered network regulates mycorrhizal symbiosis. Cell 184, 5527–5540.e18 (2021).

10. Jiang, Y. et al. Medicago AP2-Domain Transcription Factor WRI5a Is a Master Regulator of Lipid Biosynthesis and Transfer during Mycorrhizal Symbiosis. Molecular Plant 11, 1344–1359 (2018).

11. Tan, X. et al. A pair of LysM receptors mediates symbiosis and immunity discrimination in Marchantia. Cell 188, 1330–1348.e27 (2025).

12. Xue, L. et al. AP2 transcription factor CBX1 with a specific function in symbiotic exchange of nutrients in mycorrhizal Lotus japonicus. Proc Natl Acad Sci U S A 115, E9239–E9246 (2018).

13. Bravo, A., Brands, M., Wewer, V., Dörmann, P. & Harrison, M. J. Arbuscular mycorrhiza-specific enzymes FatM and RAM 2 fine-tune lipid biosynthesis to promote development of arbuscular mycorrhiza. New Phytologist 214, 1631–1645 (2017).

14. Paries, M. et al. The GRAS protein RAM1 interacts with WRI transcription factors to regulate plant genes required for arbuscule development and function. Proc. Natl. Acad. Sci. U.S.A. 122, e2427021122 (2025).

15. Jensen, P. A. Ten species comprise half of the bacteriology literature, leaving most species unstudied. 2025.01.04.631297 Preprint at 10.1101/2025.01.04.631297 (2025).

16. Stephan, T. et al. Darwinian genomics and diversity in the tree of life. Proceedings of the National Academy of Sciences 119, e2115644119 (2022).

17. Emms, D. M. & Kelly, S. OrthoFinder: solving fundamental biases in whole genome comparisons dramatically improves orthogroup inference accuracy. Genome Biol 16, 157 (2015).

18. Emms, D. M. & Kelly, S. OrthoFinder: phylogenetic orthology inference for comparative genomics. Genome Biol 20, 238 (2019).

19. Sayers, E. W. et al. GenBank 2025 update. Nucleic Acids Research 53, D56–D61 (2025).

20. UniProt Consortium. UniProt: the Universal Protein Knowledgebase in 2023. Nucleic Acids Res 51, D523–D531 (2023).

21. Mistry, J. et al. Pfam: The protein families database in 2021. Nucleic acids research 49, D412–D419 (2021).

22. Liu, F. et al. Systematic Identification, Evolution and Expression Analysis of the Zea mays PHT1 Gene Family Reveals Several New Members Involved in Root Colonization by Arbuscular Mycorrhizal Fungi. Int J Mol Sci 17, 930 (2016).

23. Blin, K. et al. antiSMASH 7.0: New and improved predictions for detection, regulation, chemical structures and visualisation. Nucleic Acids Res 51, W46–W50 (2023).

24. Huerta-Cepas, J. et al. eggNOG 5.0: A hierarchical, functionally and phylogenetically annotated orthology resource based on 5090 organisms and 2502 viruses. Nucleic Acids Res 47, D309–D314 (2019).

25. Koonin, E. V. Orthologs, Paralogs, and Evolutionary Genomics. Annual Review of Genetics 39, 309–338 (2005).

26. Greener, J. G., Kandathil, S. M., Moffat, L. & Jones, D. T. A guide to machine learning for biologists. Nat Rev Mol Cell Biol 23, 40–55 (2022).

27. He, C. et al. Identification of a de novo missense variant in the BRI3BP gene in a Holstein calf with congenital cardiac malformation and carpus valgus. Animal Genetics 56, e13494 (2025).

28. Conen, K. L., Nishimori, S., Provot, S. & Kronenberg, H. M. The transcriptional cofactor lbh regulates angiogenesis and endochondral bone formation during fetal bone development. Dev Biol 333, 348–358 (2009).

29. Charras, A. et al. P2RX7 gene variants associate with altered inflammasome assembly and reduced pyroptosis in chronic nonbacterial osteomyelitis (CNO). J Autoimmun 144, 103183 (2024).

30. Huang, H. et al. P2×7Rs: New therapeutic targets for osteoporosis. Purinergic Signal 19, 207–219 (2023).

31. Li, J., Liu, D., Ke, H. Z., Duncan, R. L. & Turner, C. H. The P2×7 nucleotide receptor mediates skeletal mechanotransduction. J Biol Chem 280, 42952–42959 (2005).

32. Xu, P., Li, X., Tang, C., Wang, T. & Xu, J. Deduction of CDC42EP3 suppress development and progression of osteosarcoma. Exp Cell Res 412, 113018 (2022).

33. Gherbi, H. et al. SymRK defines a common genetic basis for plant root endosymbioses with arbuscular mycorrhiza fungi, rhizobia, and frankiabacteria. Proc Natl Acad Sci U S A 105, 4928–4932 (2008).

34. Miyata, K. et al. OsSYMRK plays an essential role in AM symbiosis in rice (oryza sativa). Plant Cell Physiol 64, 378–391 (2023).

35. Maillet, F. et al. Fungal lipochitooligosaccharide symbiotic signals in arbuscular mycorrhiza. Nature 469, 58–63 (2011).

36. He, S. et al. Oral microbiota disorder in GC patients revealed by 2b-RAD-M. J Transl Med 21, 831 (2023).

37. Azodi, C. B., Tang, J. & Shiu, S.-H. Opening the black box: Interpretable machine learning for geneticists. Trends Genet 36, 442–455 (2020).

38. Heumos, L. et al. mlf-core: A framework for deterministic machine learning. Bioinformatics 39, btad164 (2023).

39. Pimprikar, P. et al. A CCaMK-CYCLOPS-DELLA complex activates transcription of RAM1 to regulate arbuscule branching. Curr Biol 26, 987–998 (2016).

40. Dhungel, B. A. & Govind, R. Spo0A suppresses sin locus expression in clostridioides difficile. mSphere 5, e00963–20 (2020).

41. Nan, B. & Zusman, D. R. Novel mechanisms power bacterial gliding motility. Mol Microbiol 101, 186–193 (2016).

42. Radhakrishnan, G. V. et al. An ancestral signalling pathway is conserved in intracellular symbioses-forming plant lineages. Nat Plants 6, 280–289 (2020).

43. Schober, I. et al. BacDive in 2025: The core database for prokaryotic strain data. Nucleic Acids Research 53, D748–D756 (2025).

44. Zheng, X. et al. Identification and Experimental Verification of PDK4 as a Potential Biomarker for Diagnosis and Treatment in Rheumatoid Arthritis. Mol Biotechnol (2024) doi:10.1007/s12033-024-01297-1.

45. Ramoneda, J. et al. Building a genome-based understanding of bacterial pH preferences. Sci. Adv.9, eadf8998 (2023).

46. Zhu, X. et al. Combining CRISPR–Cpf1 and Recombineering Facilitates Fast and Efficient Genome Editing in Escherichia coli. ACS Synth. Biol. (2022) doi:10.1021/acssynbio.2c00041.

